# Neural silences can be localized rapidly using noninvasive scalp EEG

**DOI:** 10.1101/2020.10.11.334987

**Authors:** Alireza Chamanzar, Marlene Behrmann, Pulkit Grover

**Author notes:** (contact: { }) (Pulkit Grover).

## Abstract

A rapid and cost-effective noninvasive tool to detect and characterize suppressed neural activity can be of significant benefit for the diagnosis and treatment of many disorders. We propose a novel algorithm, SilenceMap, for uncovering the absence of electrophysiological signals, or neural “silences”, using noninvasive scalp electroencephalography (EEG) signals. By accounting for the contributions of different sources to the power of the recorded signals, and using a novel hemispheric baseline approach and a convex spectral clustering framework, SilenceMap permits rapid detection and localization of regions of silence in the brain using a relatively small amount of EEG data. SilenceMap substantially outperformed existing source localization algorithms in estimating the center-of-mass of the silence for three pediatric patients with lobectomy, using less than 3 minutes of EEG recordings (13, 2, and 11mm vs. 25, 62, and 53mm), as well for 70 different simulated regions of silence based on a real human head model (11±0.5mm vs. 54±2.2mm). SilenceMap paves the way towards accessible early diagnosis and continuous monitoring of altered physiological properties of human cortical function.

## I. INTRODUCTION

An ongoing challenge confronting both basic scientists as well as those at the translational interface is the ability to access a rapid and cost-effective tool to uncover mechanistic details of neural function as well as the consequences of brain damage. For example, identifying the presence of stroke, establishing altered neural dynamics in traumatic brain damage, and monitoring changes in neural profile in athletes on the sidelines all pose significant hurdles. In this paper, using scalp electroencephalography (EEG) signals with relatively little data, we provide theoretical and empirical support for a novel method for the noninvasive detection of neural silences. We adopt the term “silences” or “regions of silence” to refer to the parts of brain tissue with little or no neural activity. These regions reflect ischemic, necrotic, or lesional tissue (e.g., after ischemic stroke, traumatic brain injuries (TBIs), and intracranial hematoma), resected tissue (e.g. after epilepsy surgery), or tumors [1, 2]. Dynamic regions of silence also arise in cortical spreading depolarizations (CSDs), which are slowly spreading waves of neural silences in the cerebral cortex [3–5].

Common imaging methods for detecting brain damage, e.g., magnetic resonance imaging (MRI) [6, 7], or computed tomography (CT) [8], are not portable, are not designed for continuous (or frequent) monitoring, are difficult to use in many emergency situations, and may not even be available at medical facilities in many countries. However, many medical scenarios can benefit from portable, frequent/continuous monitoring of neural silences, e.g., detecting changes in tumor or lesion size/location (e.g. expansion/shrinkage), and/or propagation of CSD in the brain. On the other hand, noninvasive scalp EEG is widely accessible in emergency situations and can even be deployed in the field with only a few limitations. It is easy and fast to setup, portable, and of lower cost compared with other imaging modalities. Additionally, unlike MRI, EEG can be recorded from patients with implanted metallic objects in their body, e.g., pacemaker [9].

One of the ongoing challenges of EEG is that of “source localization”, the process by which the location of the underlying neural activity is determined from the scalp EEG recordings. The challenge arises primarily from three issues: (i) the underdetermined nature of the problem (few sensors, many possible locations of sources); (ii) the spatial low-pass filtering effect of the distance and the layers separating the brain and the scalp; and (iii) noise, including exogenous noise, background brain activity, as well as artifacts, e.g., heart beats, eye movements, and jaw clenching [10, 11]. The localization of a region of silence, however, poses additional challenges. The most significant further challenge is in how background brain activity is treated: while it is usually grouped with noise in source localization^1^, it is of direct interest in silence localization where the goal is to distinguish normal brain activity from abnormal silences. Thus, in source localization paradigms applied to neuroscientific studies [12–14], as is also the case in the Event Related potential (ERP) paradigm [15, 16], scalp EEG signals are averaged over event-related trials to average out background brain activity and noise, permitting the extraction of the signal activity that is consistent across trials. Consequently, as we demonstrate in our experimental results below, classical source localization techniques, e.g., multiple signal classification (MUSIC) [17, 18], minimum norm estimation (MNE) [14, 19, 20], and standardized low resolution brain electromagnetic tomography (sLORETA) [21], even after appropriate modifications, fail to localize regions of silence in the brain.

In order to not average out the activity of the background sources, we estimate the contribution of each source to the recorded EEG outputs across all electrodes. This contribution is measured in an average power sense, instead of the mean, thereby avoiding canceling out the contributions of the background brain activity. Our silence localization algorithm, that we refer to as SilenceMap, estimates this contribution, and then uses tools that quantify our assumptions on the region of silence (contiguity, small size of the region of silence, and being located in only one hemisphere) to arrive at an estimate of the region of silence.

In developing an algorithm tailored to localizing silences, we observed two additional difficulties: lack of statistical models of background brain activity, and the choice of the reference electrode. The first is dealt with either by including baseline recordings (in absence of silence; which we did not have for our experimental results) or utilizing what we call a “hemispheric baseline”, i.e., an approximate equality in power measured at electrodes placed symmetrically with respect to the longitudinal fissure (see Fig. 1b). While the hemispheric baseline used here provides fairly accurate reconstructions, we note that this “baseline” is only an approximation, and an actual baseline is expected to further improve the accuracy. The second difficulty is related: to retain this hemispheric (approximate) symmetry in power, it is best to keep the reference electrode on top of the longitudinal fissure (see Fig. 1a). Using these advances, we proposed an iterative algorithm to localize the region of silence in the brain using a relatively small amount of data. Fig. 1d and e show the details of our proposed SilenceMap algorithm. In both simulation studies and real data analysis, SilenceMap outperformed existing algorithms, while using only a small amount of EEG data. Only 160s of EEG signals using 128 electrodes suffice for localizing silences in 3 participants with surgical resections for management of epileptic seizures.

**FIG. 1.**
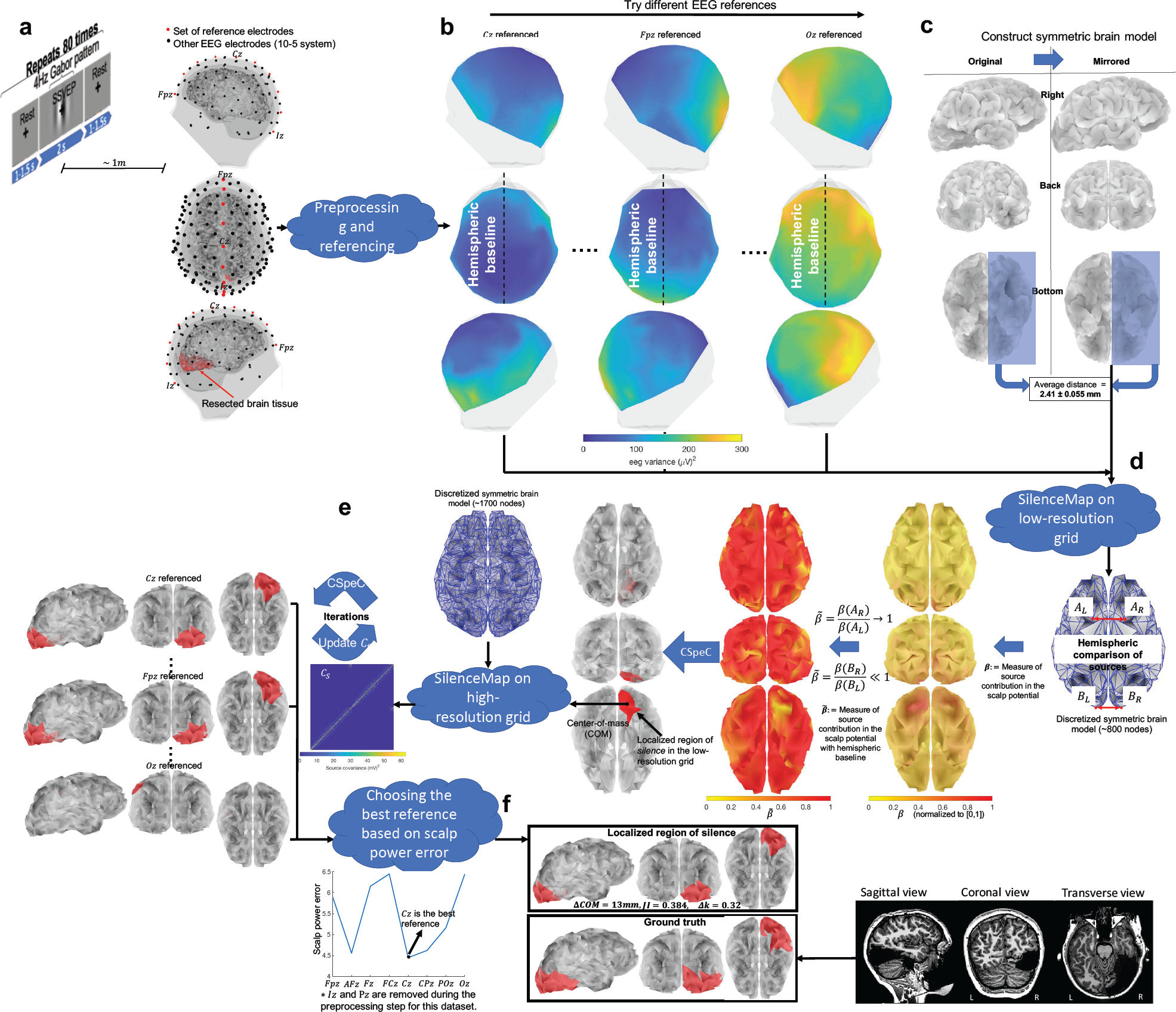
SilenceMap with baseline algorithm overview: a) The EEG recording protocol and the locations of scalp electrodes. One of 10 reference electrodes (shown in red) is chosen along the longitudinal fissure for rereferencing against. b) Average power of scalp potentials for different choices of reference electrodes. c) Symmetric brain model of a patient (UD) with right occipitotemporal lobectomy. d) Steps of the SilenceMap algorithm in a low-resolution source grid. A measure of the contribution of brain sources in the recorded scalp signals 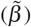 is calculated relative to a hemispheric baseline. In the brain colormap, yellow indicates no contribution. A contiguous region of silence is localized based on a convex spectral clustering (CSpeC) framework in the low-resolution grid. e) Steps of the SilenceMap algorithm in a high-resolution source grid. The source covariance matrix (*C*_*S*_) is estimated through an iterative method, and the region of silence is localized using the CSpeC framework. f) Choosing the best reference electrode to reference against (*Cz* in this example), which results in minimum scalp power mismatch (Δ*Pow*). The localized region of silence for this patient (UD) has 13mm COM distance (Δ*COM*) from the original region, with more than 38% overlap (JI = 0.384), and it is 32% smaller (Δ*k* = 0.32).

## II. RESULTS

Our SilenceMap algorithm localizes the region of silence in two steps: (1) The first step finds a contiguous region of silence in a low-resolution source grid with the assumption that sources, at this low-resolution, are uncorrelated across space. Steps of the SilenceMap algorithm in a low-resolution brain grid of a patient (UD) with right occipitotemporal lobectomy, are shown in Fig. 1d, with nodes serving as the brain sources. The sources are far enough from each other in the low-resolution grid that it maybe a reasonable approximation to assume they have independent activities (see Methods for more details). We defined a measure for the contribution of brain sources in the recorded scalp signals (*β*), i.e., the larger the *β*, the higher contribution of the brain source to the scalp potentials. However, *β* is not a perfect measure of the contribution since it is defined based on identical distribution assumption of non-silent sources in the brain, which does not hold in the real world. Therefore using *β*, as is, does not reveal the silent sources, i.e., the smallest values of *β* (yellow regions in Fig. 1d) may not be located at the region of silence. But looking closely at the bottom view of the brain reveals a significant hemispheric color difference at the region of silence (right occipito-temporal lobe). This motivated us to use a hemispheric baseline for the region of silence, i.e., instead of using *β*, we use 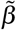 which is the ratio of *β* values of the mirrored sources, e.g., for source pair of (*A*_*L*_, *A*_*R*_) which are far from the region of silence, 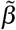 is close to 1 (red-colored sources), while for (*B*_*L*_, *B*_*R*_), where *B*_*R*_ is located in the region of silence (see Fig. 1d), this ratio is close to zero (yellow-colored sources). A contiguous region of silence is localized based on a convex spectral clustering (CSpeC) framework [22–24] in the low-resolution grid. (2) The second step of SilenceMap algorithm takes the localized region of silence in the first step as an initial guess, and through an alternating method, estimates both the source covariance matrix **C**_**s**_ and the localized contiguous region of silence in a higher resolution source grid (see Fig. 1e). In each iteration, a CSpeC framework was used to localize the region of silence based on the estimated source covariance matrix, until the center-of-mass (COM) of the localized region of silence has converged (see the Methods for more details). All steps of our silence localization algorithm are summarized in Fig. 1.

We validated the performance of our algorithm through rigorous experiments, based on simulated and real datasets. We tested the robustness of our SilenceMap algorithm with and without baseline (see Methods for more details), in different scenarios, e.g., different sizes and locations of region of silence, different EEG reference electrodes, and based on both visual and rest EEG datasets (see Fig. 1a). In both the simulated and real data, the performance of our silence localization algorithm is compared with that of the state-of-the-art source localization algorithms, namely, sLORETA, MNE, and MUSIC, which are modified appropriately for the silence localization task. We use simulations and experiments to understand how to choose the reference electrode, and what the effect of this choice is on the localization. Finally, we explored the validity of our hemispheric symmetry assumption in SilenceMap with the baseline based on a real dataset.

### Localization performance metrics

For both simulated and real experiments, we used three performance metrics for determining the accuracy of the silence localization task: (i) center-of-mass (COM) distance (Δ*COM*), (ii) Jaccard Index (JI), and (iii) size error (Δ*k*)

#### (i) COM distance

is simply defined as the Euclidean distance between the center-of-mass of the localized and actual region of silence, i.e.,

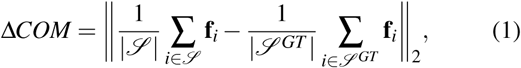

where **f**_*i*_ is the 3D location of source *i* in the brain, 𝒮 and 𝒮 ^*GT*^ are the set of source indices of the localized region of silence and its ground truth, respectively. Δ*COM* basically measures how far the localized region of silence is from the ground truth.

#### (ii)JI

was first defined by Jaccard in [25]. It is a widely used performance measure for the 2D image segmentation tasks [26]. In the silence localization task, since we are segmenting the region of silence in 3D space, we can calculate the JI based on the nodes/sources in the discretized brain as follows:

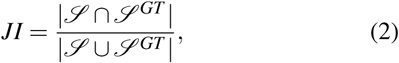

which measures how well the localized region of silence overlaps with the ground truth region in the brain, and it assumes values between 0 (no overlap) and 1 (perfect overlap). If there is minimal overlap and/or there is a large mismatch between the size of these two regions, *JI* has a small value.

#### (iii) Size error

measures the error in estimation of the size of the region of silence, and is simply defined as follows:

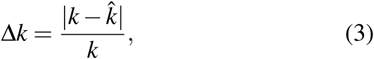

where 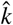 is the estimated number of silent sources in the localized region of silence.

### Simulations

We simulated the scalp EEG recordings of regions of silence in the brain, following the assumptions we made in Section IV A, and tested the performance of the SilenceMap algorithm proposed in this paper.

#### Simulation result

We simulated scalp differential recordings for 70 different regions of silence, with the size of *k* = 50, on a high-resolution source grid with *p* = 1744 sources, and at varying locations on the cortex. The simulated regions of silence lie in only one hemisphere (see the assumptions in Section IV A), and they are located no deeper than 3cm from the surface of scalp, which covers the entire thickness of the gray matter [27–29], while excluding deep sources located in the longitudinal fissure. In the longitudinal fissure, the source dipoles are located deep inside the brain, and mostly oriented tangential to the surface of the scalp, which makes it hard for the EEG to record their electrical activity [11]. The non-silent sources are assumed to have an identical distribution and correlation across space. The detailed steps for the simulation are available in the Methods (see Section IV F). For the SilenceMap algorithm, we tried the 10 different reference electrodes located along the longitudinal fissure, i.e., *F pz, AFz, Fz, FCz, Cz, CPz, Pz, POz, Oz*, and *Iz*, and choose the one with the minimum power mismatch Δ*Pow* defined in (39). We reported the performance measures, i.e., Δ*COM, JI*, and Δ*k* under the average signal-to-noise ratio (*SNR*_*avg*_) level of 9dB (see the Methods for definition of *SNR*_*avg*_). For the SilenceMap algorithm, we reported the convergence rate (*CR*) as well, which is the ratio of the number of converged cases over the total number of simulated regions of silence in the simulation experiment. The results of the simulations are summarized in Table I, where Δ*COM*, JI, and Δ*k* are reported in the format of “mean ± Standard Error (SE)”. Based on the results, our proposed SilenceMap algorithm outperformed the state-of-the-art source localization algorithms: it has 43mm smaller average COM distance, 46% more average overlap (JI), and 252% smaller size error, compared to the best performance among the modified source localization algorithms. The simulated dataset is based the identical distribution assumption of brain sources (see Section IV F). However, this assumption does not appear to hold in the real dataset, where SilenceMap *with* baseline performs significantly better compared to SilenceMap without baseline (see Fig. 3). The list of all parameters and their values we have used in the implementation of the SilenceMap algorithm is available in Appendix X D.

**TABLE I.**
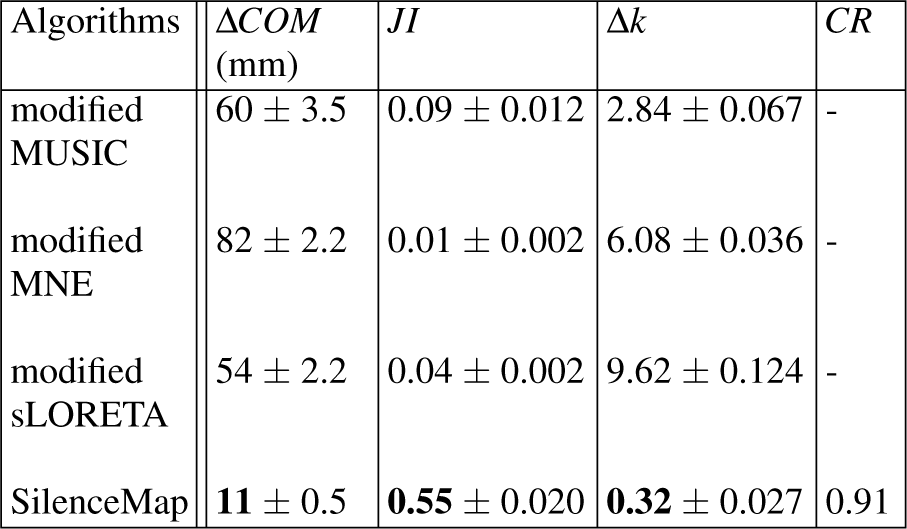
Simulation experiment results (*SNR*_*avg*_ = 9*dB, k* = 50)

**FIG. 2.**
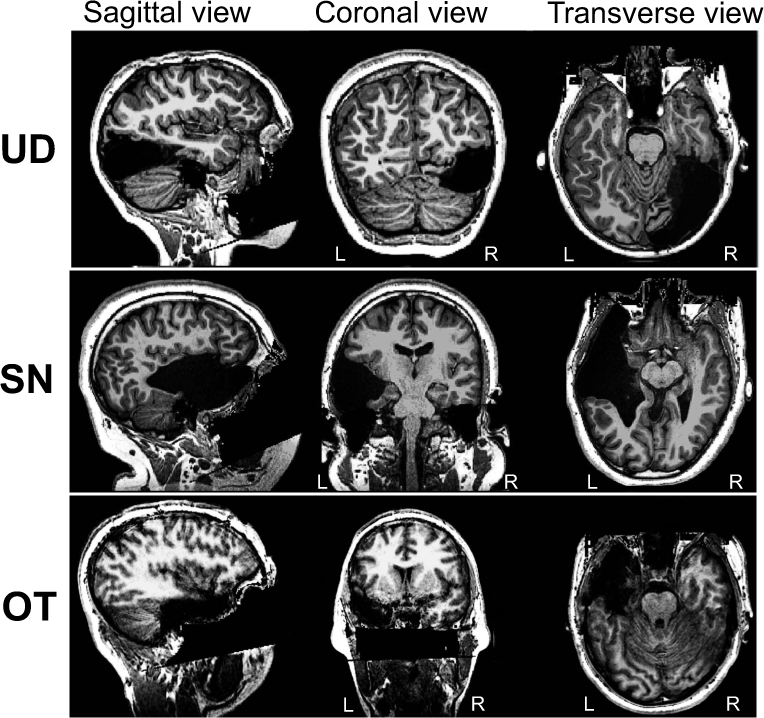
Structural MRI scans of the three participants in the real dataset: UD with right occipitotemporal lobectomy, SN and OT with left temporal lobectomy. We have stripped away facial features to ensure anonymity of participants.

**FIG. 3.**
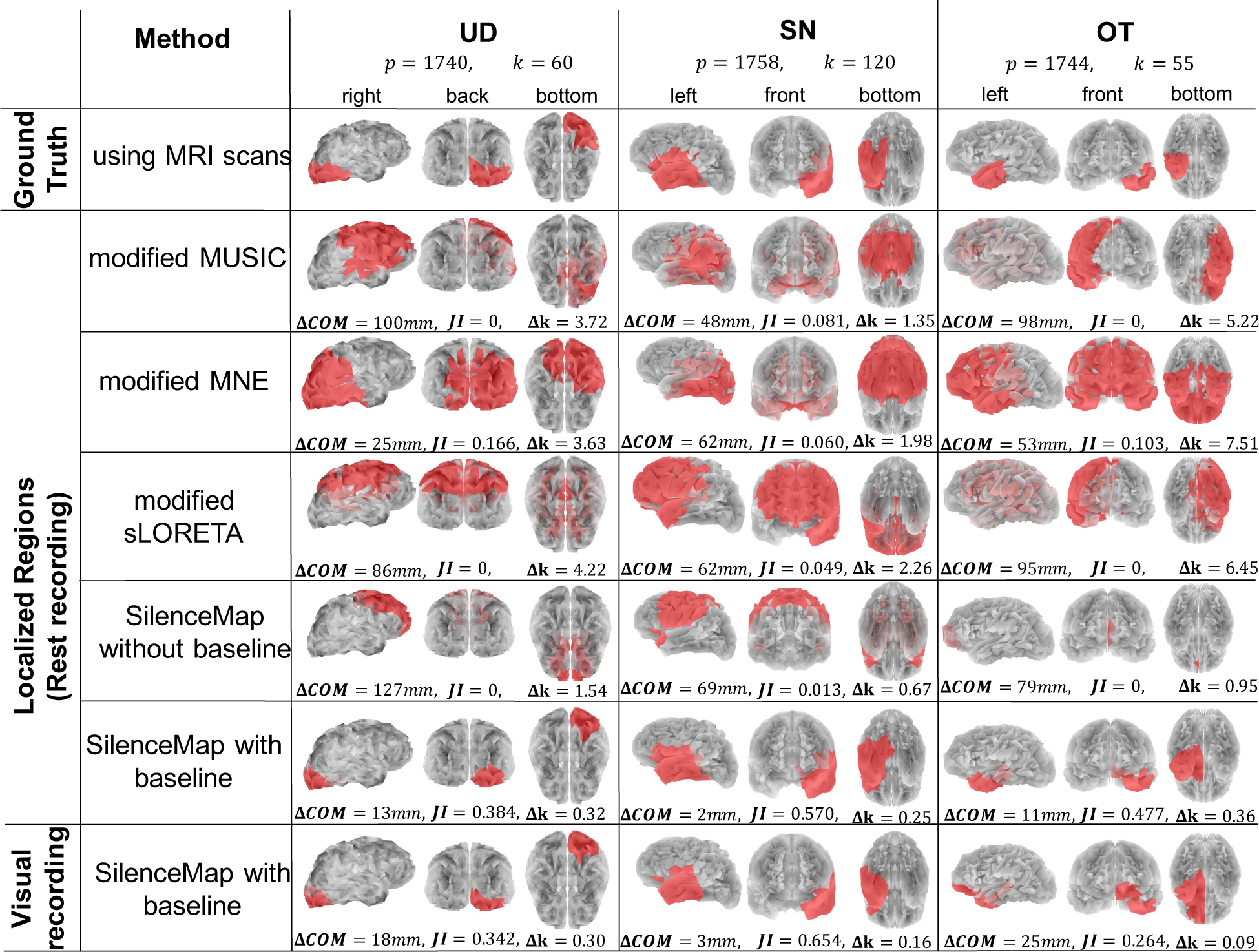
Performance of SilenceMap on a real EEG dataset: the first row shows the extracted ground truth regions of silence (red regions) overlaid on the resected cortical region of three patients based on their symmetric brain models extracted from the structural MRIs (see the MRI scans in Fig. 2); the second, third, and fourth rows show the performance in localization of the silent region using modified source localization algorithms (MNE, MUSIC, and sLORETA), through both visual illustration (red regions) and using performance metrics of center-of-mass (COM) distance (Δ*COM*), Jaccard Index (*JI*), and size error (Δ*k*). The fifth row shows the performance of our SilenceMap algorithm without baseline, and the last two rows show the localization performance of our SilenceMap algorithm with baseline, based on the Rest and Visual recordings respectively. *p* is the total number of sources in each brain model, and *k* is the size of ground truth region of silence.

### Comparison of SilenceMap, with and without baseline, with source localization algorithms

We tested the performance of our silence localization algorithm under different simulated scenarios, as well as real experiments, and compare its performance with the state-of-the-art source localization algorithms, modified for the silence localization task. We have explained the details of modified MNE, MUSIC, and sLORETA algorithms for the silence localization task in Methods. For all of the modified source localization methods, we choose *C*_*z*_ scalp electrode as the reference electrode. We considered the effect of the reference electrode in the modified MNE, MUSIC, and sLORETA, to have a fair comparison with our proposed SilenceMap algorithm (see Methods). Based on the simulation results in Table I, among the modified source localization algorithms, sLORETA shows the minimum average COM distance of 54mm, and MUSIC shows the maximum average overlap of 9% (JI = 0.09), and the minimum average size error of 284% (Δ*k* = 2.84). This performance is still poor for the silence localization task, while our SilenceMap algorithm shows a good performance based on the simulation results in Table I (Δ*COM* = 11mm, JI = 0.55, Δ*k* = 0.32). Based on these results, source localization algorithms, even after proper modifications, perform poorly in localizing the regions of silence in the brain.

### Real Data

We tested the performance of our silence localization algorithm, and compared it with the modified source localization algorithms, based on a real dataset of patients who have undergone lobectomy surgery, and have a clearly-defined resected region in their brain. It is this region of silence that we intend to localize.

#### Dataset

We recorded EEG signals using a BioSemi ActiveTwo system (BioSemi, Amsterdam), with a sampling frequency of 512 *Hz*, using a 128-electrode cap with electrodes located based on the standard 10-5 system [30]. In addition, we used four electrodes around the eyes, specifically, a pair on the top and bottom of the right eye to detect the vertical eye movements and blinks, and a pair at the outer canthi of each eye to monitor horizontal eye movements. One electrode was placed on the left collar bone to monitor the heart beats, and two electrodes were placed on the mastoids. All electrodes were differentially recorded relative to the standard common-mode-sense (CMS) and driven-right-leg (DRL) electrodes. During the acquisition of EEG data, the participant viewed a screen, located roughly 1m away. A grating pattern of black and white bars was displayed at the center of the display along with a fixation cross for 2 seconds, followed by a rest state of 1-1.5 seconds, where a fixation cross was displayed on a gray-colored background (see Fig. 1a). We repeated this sequence 80 times during the recording session. We used the Rest and Visual sections of the recorded signal separately for the localization and compared the results from these analyses in Fig. 3. The steps for data analyses and pre-processing are available in the Methods, part IV E.

### Participants

Three male pediatric patients were recruited for this experiment. Two patients (SN and OT) had resections in the left hemisphere and one (UD) had a resection in the right hemisphere. In two of these patients (OT and UD), lobectomy surgery was performed to control pharmacoresistant epilepsy, and in the third patient (SN) surgery was performed for an emergent evacuation of cerebral hematoma at day one of life. More information about these patients is included in Table II.

**TABLE II.**
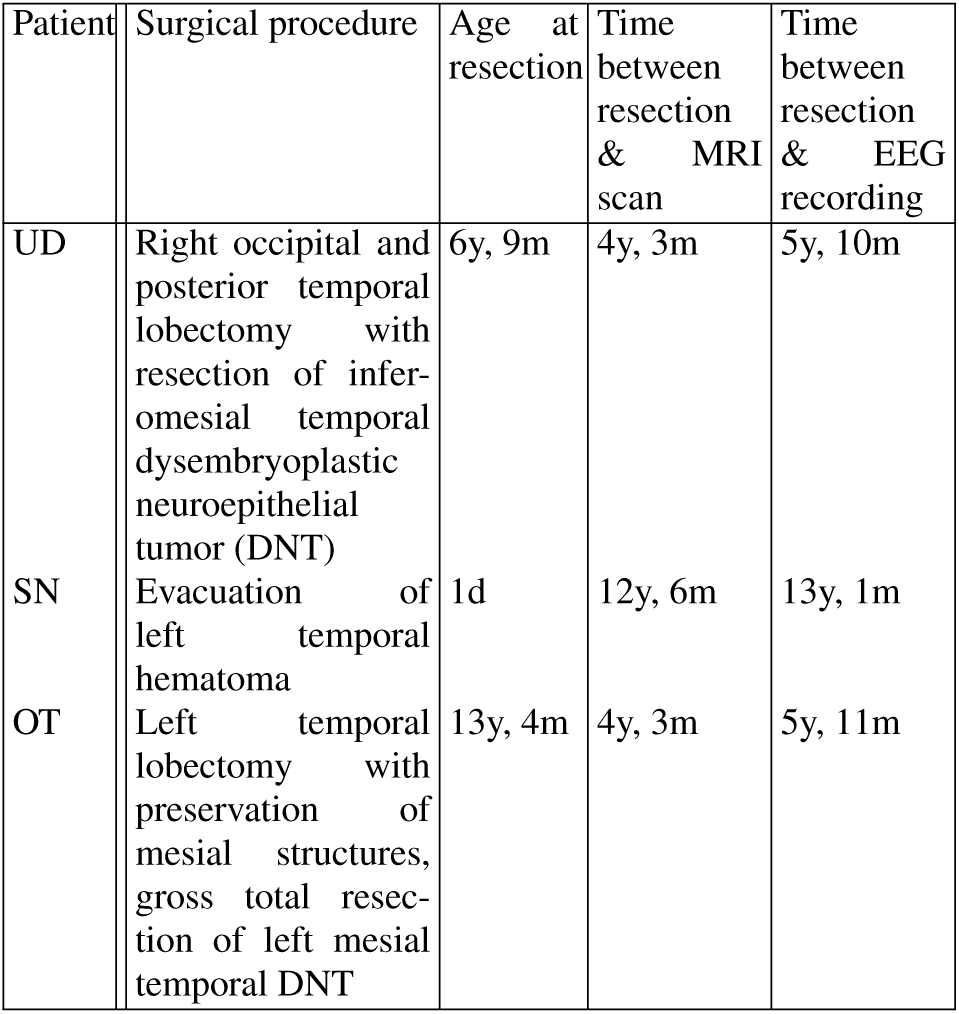
Surgery history of patients [31].

OT signed a consent form, and the parents of SN and UD consented to their participation as they were minors. They both provided assent. All procedures were approved by the Carnegie Mellon University Institutional Review Board (IRB). The MRI scans of these participants are shown in Fig. 2, where the resected sections can be seen as large asymmetric dark regions [31–33]. The ground truth regions of silence are extracted based on these MRI scans (see Methods for more details). First row in Fig. 3 shows the extracted ground truth regions of silence in the symmetric brain models of the three participants. The intact hemisphere is mirrored across the longitudinal fissure to construct these brain models (see Fig. 1c and Methods for more details). These patients have different sizes of regions of silence: UD has a region of silence with *k* = 60 nodes out of total *p* = 1740 nodes in the brain, SN has *k* = 120 out of *p* = 1758 nodes, and OT has *k* = 55 out of *p* = 1744 total nodes.

#### Results of real database

We applied the SilenceMap algorithm, along with the modified source localization algorithms, i.e., MNE, MUSIC, and sLORETA, on the preprocessed EEG recordings of the three participants in the real dataset, and the performance of silence localization is calculated based on the extracted ground truth regions from the post-surgery MRI scans of these patients (see Fig. 2). The visual illustration of localized regions of silence (shown in red color on the gray-colored semi-transparent brains), along with their ground truth regions and their corresponding performance measures are all shown in Fig. 3. Based on the Rest dataset, our SilenceMap algorithm with hemispheric baseline outperforms the modified source localization algorithm: it reduces the COM distance by 12mm, 46mm, and 42mm for UD, SN, and OT respectively, compared to the best performance among the source localization algorithms. It also improves the overlap (JI) by 22%, 49%, and 37%, and the size estimation by 122%, 42%, and 59% for UD, SN, and OT respectively. The SilenceMap algorithm with baseline performs well with values of Δ*COM* = 2*mm, JI* = 0.570, and Δ*k* = 0.25 based on the Rest set, and Δ*COM* = 3*mm, JI* = 0.654, and Δ*k* = 0.09 based on the Visual set. Comparing the results of Visual and Rest datasets for SilenceMap with baseline shows that, as expected, the localized regions of silence remain largely the same. This suggests that for each participant with a specific region of silence in the brain tissue, the algorithm can localize the region, regardless of the type of the task performed (Visual or Rest) by the participants during the EEG recording. In SilenceMap with baseline, based on the minimum value of the power mismatch (Δ*Pow* defined in (39) in the Methods), the best reference electrodes for UD, SN, and OT were found as *Cz, Cz*, and *CPz* respectively, for the Rest set, and *Cz, Pz*, and *CPz* for the Visual set. Based on the results of the Visual set, participants OT shows the poorest localization performance, which might be due to the consistent jaw clenching the participant had during the recording session which affected the EEG signals, even after appropriate preprocessing steps. Jaw clenching is recognized as one of the most severe artifacts in EEG recording which adversely impacts the signals of most EEG electrodes [34].

Unlike the simulation results, without baseline the SilenceMap algorithm failed to localize the region of silence based on the real dataset. As mentioned before, one explanation for this is the assumption of the identical distribution of sources in designing the algorithm, which does not hold in the real data and we need to use the hemispheric baseline to be able to localize the region of silence.

### Validity of hemispheric symmetry assumption in SilenceMap with baseline

The hemispheric baseline approach used in the SilenceMap algorithm is based on an approximate hemispheric symmetry assumption of the brain source activities in the healthy parts of the brain. To further explore the validity of this assumption, we quantified this hemispheric symmetry based on the scalp average power of a neurologically healthy control subject (DH, male, 25yr) whose EEG data were collected using the same protocol as that used for the patients (see Fig. 1a for the EEG recording protocol). DH signed a consent form. Excluding the 10 electrodes on the longitudinal fissure (red electrodes in Fig. 1a), we calculated the mean absolute difference (*MAD*) of average power of pairs of scalp electrodes which are symmetric with respect to the longitudinal fissure, e.g., (*C*1,*C*2), (*T* 7,*T* 8), and so on, as follows:

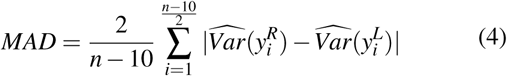

where, 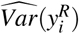 is the estimated variance of the recorded EEG signals referenced to the *Cz* electrode, preprocessed, and denoised signal 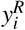 at the electrode *i* on the right hemisphere (see Methods for noise removal steps), and 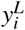 is the signal of the corresponding electrode on the left hemisphere, and *n* = 128 is the total number of electrodes. Based on Fig. 4, *MAD* for the control subject is calculated based on the Rest set as 4.1 (*µV*)^2^, while for the UD, SN, and OT patients with regions of silence *MAD* is 23.3, 14.2, 16.5 (*µV*)^2^ respectively. The control subject had significantly smaller hemispheric difference of scalp power compared to the patients with regions of silence in their brain. This result supports the fact that using the hemispheric baseline is helpful in localization of regions of silence, which are located in either the left or right hemispheres. There are two main reasons that the *MAD* of the healthy control is not perfectly symmetric: (i) the sources in brain do not have perfectly symmetric activities and brain sources have non-identical brain activities, (ii) the structure of brain and the head (scalp, skull, CSF, and brain) is not perfectly symmetric, which results into a non-symmetric reflection/transformation of brain activities to the scalp potentials. The second issue is addressed in the SilenceMap algorithm, by normalization of the measure of source contribution (*β* in equation (35), in the Methods) based on the head structure asymmetry. One possible direction to improve the performance of the SilenceMap algorithm is to take into account the non-identical distribution of sources in brain (and perhaps use a more realistic model for the source covariance matrix *C*_*S*_) and normalize the source contribution measure accordingly.

**FIG. 4.**
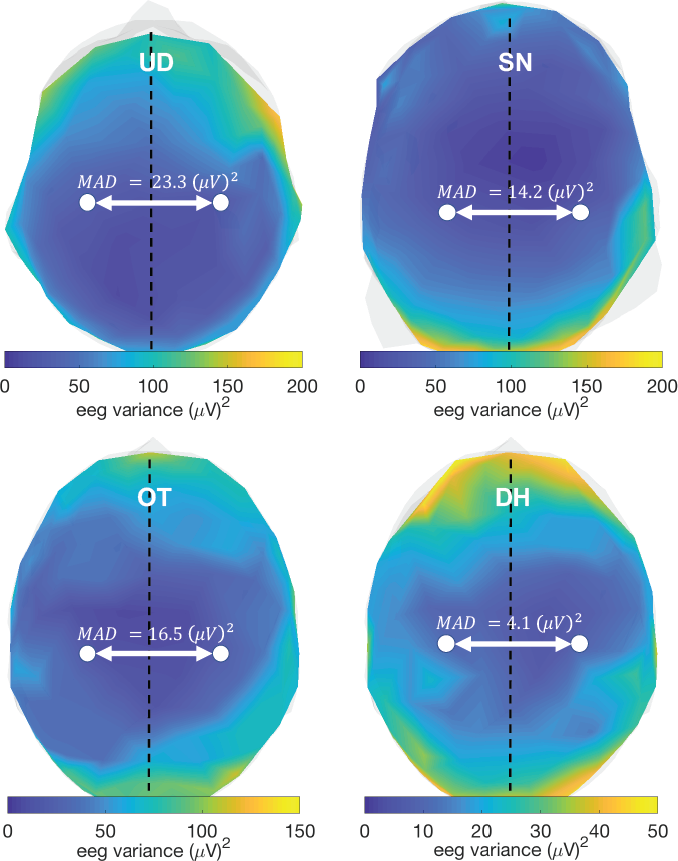
Quantification of hemispheric symmetry of scalp average power in a healthy control subject (DH), in comparison to the three patients who have resected brain regions (UD, SN, and OT). Mean absolute difference of scalp average power (*MAD*) is reported for each subject. The control subject shows significantly smaller *MAD* compared to the three patients with cortical regions of silence.

## III. DISCUSSION

SilenceMap with baseline successfully localized the region of silence of three patients who had resected brain tissue with different sizes and locations, based on only 160s of scalp EEG recordings. As the first silence localization algorithm, SilenceMap significantly outperformed the state-of-the-art source localization algorithms, which are modified for the silence localization task, and reduces the distance error (Δ*COM*) up to 46mm. Source localization algorithms, even after appropriate modifications, failed to localize the region of silence, because the background brain activity is usually grouped with noise in the source localization task, and in most of the source localization algorithms this component is averaged out across event-related trials, while including the background brain activity is a crucial component for the silence localization task.

### SilenceMap can localize the regions of silence with a tiny amount of EEG data

As we showed in Section II, our SilenceMap algorithm successfully localized the regions of silence based on only 160s of EEG data. Although this is already considered as a small amount of data, we were wondering how our algorithm performs if we reduce the temporal length of the data used for the silence localization task. To find the answer, we did a grid-search for the length of the EEG signals in the interval of [20, 40, 80, 120, 160]s, and for each temporal length we quantified the localization performance of SilenceMap. Based on the results, for the participant UD, using only 80s of data showed almost the same localization performance as 160s (Δ*COM* = 17*mm, JI* = 0.382, Δ*k* = 0.30), while 40s of data and less showed significant reduction in the localization performance. For participant SN, the minimum possible amount of data, without compromising the localization performance, is only 40s (Δ*COM* = 9*mm, JI* = 0.440, Δ*k* = 0.20), while for participant OT, any amount of data less than 160s showed reduction in the performance of silence localization. This observation might be due to the noisy EEG recording of OT, as mentioned in Section II. These sets of results suggest that in case of having a good EEG recording (without significant noise and artifacts), SilenceMap with baseline is able to localize the regions of silence, with only a tiny amount of data. This paves the way towards designing a monitoring system for the moving regions of silence in the brain such as expanding tumors and lesions, and even CSD waves based on our SilenceMap algorithm with some modifications.

### Introduced error in the silence localization by using the symmetric brain models

In this paper, we used the symmetric brain models of the patients with lobectomy, since the pre-surgery MRI scans of these patients were not available (and these may not even have been symmetrical in the first instance). Fig. 5 shows the symmetric brain models of UD, SN, and OT, along with their original brain models, which have resected regions. To quantify the introduced error in the silence localization by using the symmetric brain, instead of the original brain model, we calculated the average distance of sources/nodes of the intact part of the hemisphere with a missing section to the corresponding sources/nodes of the other hemisphere (the structurally preserved hemisphere) mirrored across the longitudinal fissure (see Fig. 5). Following the 3D shape matching approach in [35], for a specific source/node in the brain hemisphere with the region of silence, the corresponding source in the mirrored hemisphere is defined as the node with the minimum distance to that specific source. Based on the results, the defined average distance between the symmetric brain model and the original brain model is 2.41±0.055mm, 2.50±0.043mm, and 2.03±0.044mm, for UD, SN, and OT, respectively. We excluded the resected parts of the brain in calculating the average distance between the symmetric brain model and the original brain model in UD, SN, and OT. To make sure this average distance is not affected by this exclusion of the resected regions, we also calculated this hemispheric distance in three healthy controls OAS1 0004 MR1 (male, 28yr), OAS1 0005 MR1 (male, 18yr), and OAS1 0034 MR1 (male, 51yr), where there are no resected parts (see Fig. 5). We used an open source MRI database (OASIS-1^2^ [36]) to obtain real brain models of these three healthy controls. Based on the results, the average distance between the symmetric brain model and the original brain model was 2.33±0.012mm, 2.78±0.016mm, and 2.35±0.012mm, for OAS1 0004 MR1, OAS1 0005 MR1, and OAS1 0034 MR1, respectively. In fMRI studies, an acceptable motion and voxel displacement is usually up to 3mm [37], and since the average distance of the symmetric and the original brain models is less than 3mm, using the symmetric brain model seems to be a reasonable choice for the silence localization.

**FIG. 5.**
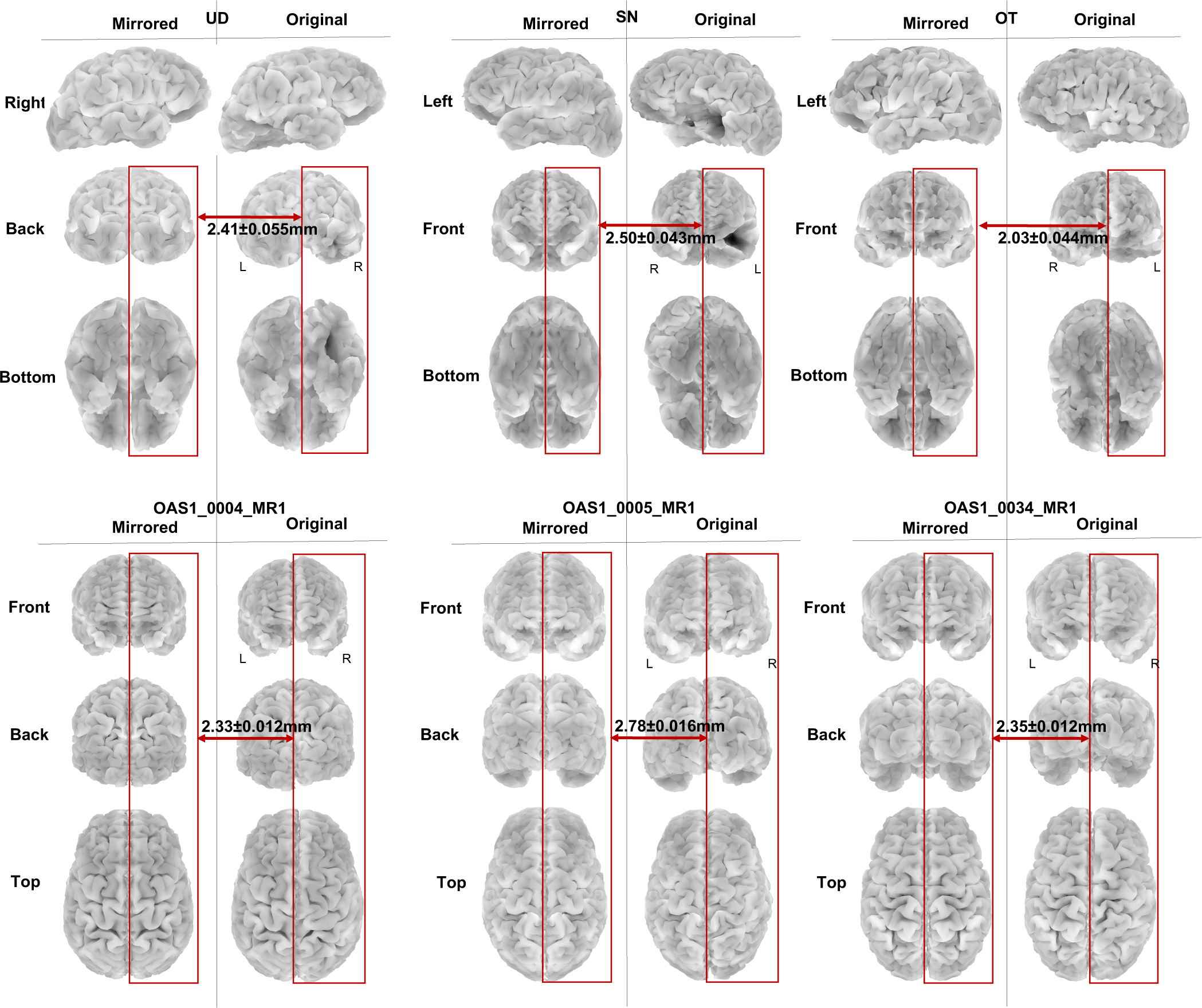
Average distance of the symmetric brain model from the original brain model in patients with lobectomy, i.e., UD with right occipitotemporal lobectomy, SN with left temporal lobectomy, and OT with left frontotemporal lobectomy, as well as in three healthy controls, i.e., OAS1 0004 MR1, OAS1 0005 MR1, and OAS1 0034 MR1. The average distance for all patients and healthy controls are less than 3mm, which makes the symmetric brain model a reasonable choice for the silence localization task, as it does not introduce significant error.

### Limitations and future directions

SilenceMap has its own limitations and shortcomings, which can serve as the focus of future investigations: (i) as mentioned in Methods, Section IV A, for simplicity, SilenceMap assumes that there is one region of silence in the brain and it is located in only one hemisphere, as is the case for the individuals in our real and simulated dataset. One can modify the SilenceMap algorithm so that it can localize multiple regions of silence in the brain, including even defective regions that span both hemispheres. (ii) Based on the results, the SilenceMap algorithm showed significant improvement in estimation of size of region of silence, compared to the state-of-the-art modified source localization algorithms. This size estimation in the SilenceMap algorithm is based on the minimization of scalp average power error, as defined in equation (27) in Methods, which shows an average error of about 30% based on the real dataset in our paper. For those applications where there is a need for more precise estimation of the size of the regions of silence, SilenceMap needs to be improved. (iii) Finally, the proposed silence localization algorithm is designed to localize the stationary (or structural) regions of silence. Designing an algorithm to track and localize the moving or functional regions of silence, such as CSD propagation across the cortex, is a future direction in our silence localization research.

## IV. METHODS

### A. Notation and Problem Statement

#### Notation

In this paper, we use non-bold letters and symbols (e.g., *a, γ*, and *θ*) to denote scalars; lowercase bold letters and symbols (e.g., **a, *γ***, and ***θ***) to denote vectors; uppercase bold letters and symbols (e.g., **A, E**, and **Δ**) to denote matrices, and script fonts (e.g., 𝒮) to denote sets.

#### Problem statement

Following the standard approach in the source localization problems, we use the linear approximation of the well-known Poisson’s equation to write a linear equation, which relates the neural electrical activities in the brain to the resulting scalp potentials [38, 39]. This linear equation is called “forward model” [40]. In this model, each group of neurons are modeled by a current source or dipole, which is assumed to be oriented normal to the cortical surface [14].

The linear forward model can be written as below:

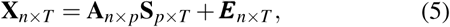

where **A** is the forward matrix, **X** is the matrix of measurements where each row represents the potentials recorded at one electrode, with reference at infinity, across time. **S** is the matrix of source signals, ***E*** is the measurement noise, *T* is the number of time points, *p* is the number of sources, and *n* is the number of scalp sensors.

In practice, we do not have the matrix **X**, since the reference at infinity cannot be recorded. Only a differential recording of scalp potentials is possible. If we define a (*n* − 1) × *n* matrix **M** with the last column to be all −1 and the first *n* − 1 columns compose an identity matrix, the differential scalp signals, with the last electrode’s signal as the reference, can be written as follows:

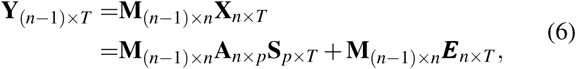

where **Y** is the matrix of differential signals of scalp, **M** is a matrix, which transforms the scalp signals with reference at infinity in the matrix **X** to the differential signals in **Y**. Equation (6) can be rewritten as follows:

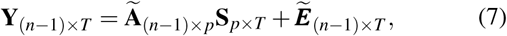

where 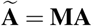, and 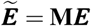.

##### Objective

Given **M, Y** and **A**, estimate the region of silence in **S**.

For this objective, we consider two different scenarios: (1) there are no baseline recordings for the region of silence, i.e., no scalp EEG recording is available where there is no region of silence, (2) with baseline recording, i.e., we consider the recording of the hemisphere of the brain, left or right, which does not have any region of silence, as a baseline for the silence localization task.

We make the following assumptions: (i) **A** and **M** are known, and **Y** has been recorded. (ii) 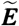 is additive white noise, whose elements are assumed to be independent across space. Thus at each time point, the covariance matrix is **C**_*z*_ given by:

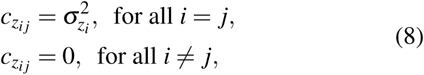

where 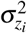 is the noise variance at electrode *i*, and it is assumed to be known (to see how this might be estimated see Section X B). (iii) *k* rows of **S** correspond to the region of silence which are rows of all zeros. The correlations of source activities reduces as the spatial distance between the sources increases. We assume a spatial exponential decay profile for the source covariance matrix **C**_*s*_, with identical variances for all non-silent sources 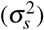:

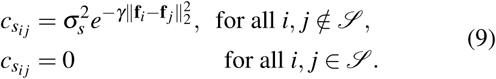

where **f**_*i*_ is the 3D location of source *i* in the brain, *γ* is the exponential decay coefficient, and 𝒮 is the set of indices of silent sources (𝒮 := *{i*| *s*_*it*_ = 0 for all *t* ∈ *{*1, 2, *T}}*). We assume that the elements of **S** have zero mean, and follow a *WSS* process. (iv) **M** is a (*n* − 1)× *n* matrix where the last column is −𝟙 _*n*− 1× 1_ and the first *n*− 1 columns form an identity matrix (**I**_(*n*− 1) × (*n*− 1)_). (v) We assume *p*− *k* » *k*, where *p*− *k* is the number of active, i.e., non-silent sources, and *k* is the number of silent sources. (vi) Silent sources are contiguous. We define contiguity based on a *z*-nearest neighbor graph, where the nodes are the brain sources (i.e., vertices in the discretized brain model). In this *z*-nearest neighbor graph, two nodes are connected with an edge, if either or both of these nodes is among the *z*-nearest neighbors of the other node, where *z* is a known parameter (see Section IV to see what values of *z* can be used). A contiguous region is defined as any connected subgraph of the defined nearest neighbor graph, i.e., between each two nodes in the contiguous region, there is at least one connecting path. (vii) For simplicity, we assume that silence lies in only one hemisphere (as is the case for the 3 individuals examined in the Results).

##### With baseline recordings

In the absence of baseline recordings, estimating the region of silence proves difficult. In order to exploit prior knowledge about neural activity, we use the approximate symmetry of power of neural activity in the two hemispheres of a healthy brain (see the Results for more details on the hemispheric symmetry of scalp potentials, along with examples from the real dataset). (viii) As an additional simplification, we assume that even when there is a region of silence, if the electrode is located far away from the region of silence, then the symmetry still holds. E.g., if the silence is in the occipital region, then the power of the signal at the electrodes in the frontal region (after subtracting noise power) is assumed to be symmetric in the two hemispheres (mirror imaged along the longitudinal fissure). This is only an approximation because (a) the brain activity is not completely symmetric, and (b) a silent source affects the signal everywhere, even far from the silent source (see Fig. 4 in the Results). Nevertheless, as we will see, this assumption enables more accurate inferences about the location of the silence region in real data using the SilenceMap algorithm with baseline, in comparison to the SilenceMap without baseline.

We first explain the details of this two-step algorithm under the condition where we do not have any baseline in Section IV B, and then under the condition where we consider a hemispheric baseline in Section IV C.

### B. SilenceMap without baseline recordings

If we do not have any baseline recording, we design the two-step silence localization algorithm as follows:

#### Low-resolution grid and uncorrelated sources

For the iterative method in the second step, we need an initial estimate of the region of silence to select the electrodes whose powers are affected the least by the region of silence. We coarsely discretize the cortex to create a very low resolution source grid with sources that are located far enough from each other, so that the elements of **S** can be assumed to be uncorrelated across space:

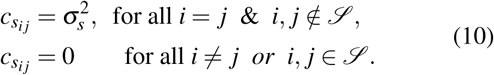

Under this assumption of uncorrelatedness and identical distribution of brain sources in this low resolution grid, we find a contiguous region of silence through the following steps:

##### (i) Cross-correlation

Equation (7) can be written in the form of linear combination of columns of matrix **Ã**as follows:

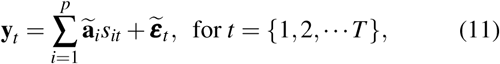

where *s*_*it*_ is the *i*^*th*^ element of the *t*^*th*^ column in **S, Y** = [**y**_1_, *…*, **y**_*T*_] ∈ ℝ^(*n*−1)*×T*^, **S** = [**s**_1_, *…*, **s**_*T*_] ∈ ℝ^*p*×*T*^, **Ã** = [**ã**_1_, *…*, **ã**_*p*_] ∈ ℝ^(*n*−1)×*p*^, and 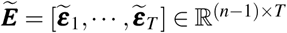

Based on equation (11), each column vector of differential signals, i.e., **y**_*t*_, is a weighted linear combination of columns of matrix **Ã**, with weights equal to the corresponding source values. However, in the presence of silences, the columns of **Ã** corresponding to the silent sources do not contribute to this linear combination. Therefore, we calculate the cross-correlation coefficient *µ*_*qt*_, which is a measure, albeit an imperfect one^3^, of the contribution of the *q*^*th*^ brain source to the measurement vector **y**_*t*_ (across all electrodes) at the *t*^*th*^ time-instant, defined as follows:

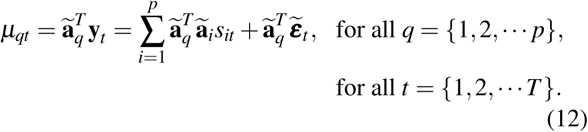

##### (ii) Estimation of variance of µ_*qt*_

In this step, we estimate the variances of the correlation coefficients calculated in the step *(i)*. Based on equation (12) we have:

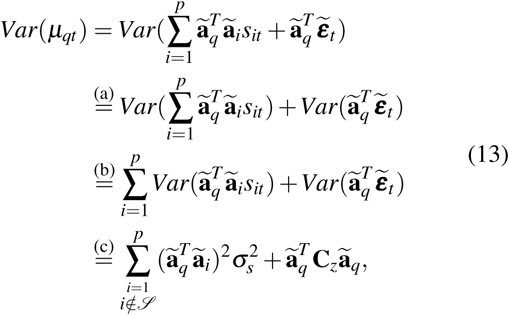

where 𝒮 is the indices of silent sources. In (13), the equality (a) holds because of independence of noise and sources, and the assumption that they have zero mean, (b) holds because the elements of **S**, i.e., *s*_*it*_’s, are assumed to be uncorrelated and have zero mean in the low resolution grid, and (c) holds because *s*_*it*_’s are assumed to be identically distributed. It is important to note that 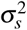 in (13) is a function of source grid discretization and it does not have the same value in the low-resolution and high-resolution grids. We estimate the variance of *µ*_*qt*_ using its power spectral density, as is explained in detail in the Appendix X A.

Based on equation (13), the variance of *µ*_*qt*_, excluding the noise variance, can be written as follows:

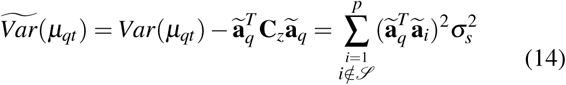

where 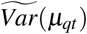 is the variance of *µ*_*qt*_ without the measurement noise term, which is a function of the size and location of region of silence through the indices in 𝒮. However, this variance term, as is, cannot be used to detect the silent sources, since some sources may be deep, and/or oriented in a way that they have weaker representation in the recorded signal **y**_*i*_, and consequently have smaller Var (*µ*_*qt*_) and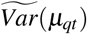.

##### (iii) Source contribution measure (***β*** )

To be able to detect the silent sources and distinguish them from sources which inherently have different values of 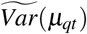, we need to normalize this variance term for each source by its maximum possible value, i.e., when there is no silent source 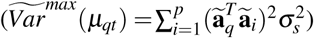

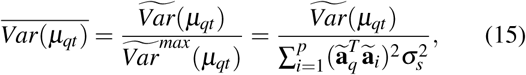

where 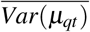 is the normalized variance of *µ*_*qt*_, without noise, and it takes values between 0 (all sources silent) and 1 (no silent source). Note that it does not only depend on whether *q* ∈ 𝒮, where 𝒮 is the set of indices of silent sources. In general, this normalization requires estimation of source variance 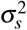, but under the assumption that sources have identical distribution, they all have identical variances. Therefore, 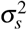 in the denominator of (15) is the same for all sources. We multiply both sides of (15) by 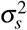 and obtain:

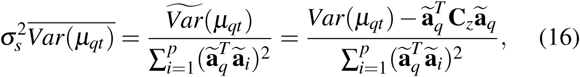

Therfore,

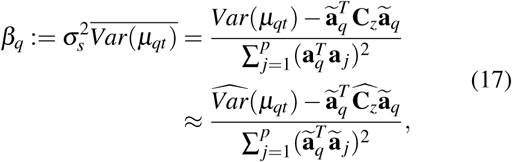

where *β*_*q*_ is called the contribution of the *q*^*th*^ source in the differential scalp signals in **Y**, which takes values between 0 (all sources silent) and 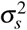 (no silent sources). In (17), 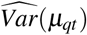 is an estimate of variance of *µ*_*qt*_, and 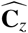 is an estimate of noise covariance matrix (see Appendix X A and X B to see how these estimates might be obtained).

##### (iv) Localization of silent sources in the low-resolution grid

In this step, we find the silent sources based on the *β*_*q*_ values defined in the previous step, through a convex spectral clustering (CSpeC) framework as follows:

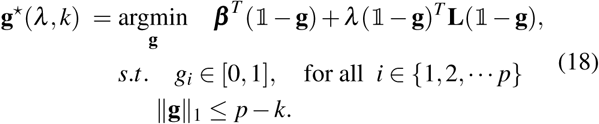

where ***β*** ^*T*^ = [*β*_1_, *…, β*_*p*_] is the vector of source contribution measures, **g** = [*g*_1_,, *g*_*p*_]^*T*^ is a relaxed indicator vector with values between 0 (for silent sources) and 1 (for active sources), *k* is the number of silent sources, i.e., the size of the region of silence, *λ* is a regularization parameter, and **L** is a graph Laplacian matrix defined in (23) below. Based on the linear term in the cost function of (18), the optimizer finds the solution **g**^*^ that (ideally) has small values for the silent sources, and large values for the non-silent sources. The *C*1 norm convex constraint controls the size of region of silence in the solution. To make the localized region of silence contiguous, we have to penalize the sources which are located far from each other. This is done using the quadratic term in the cost function in (18) and through a graph spectral clustering approach, namely, relaxed RatioCut partitioning [22–24]. We define a *z*-nearest neighbor undirected graph with the nodes to be the locations of the brain sources (i.e., vertices in the discretized brain model), and a weight matrix **W** defined as follows:

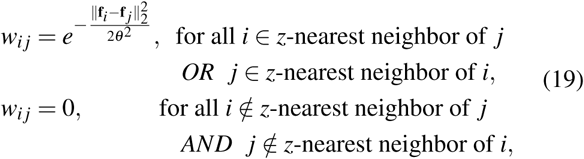

where the link weight is zero (no link) between node *i* and *j*, if node *i* is not among the *z*-nearest neighbors of *j*, and node *j* is not among the *z*-nearest neighbors of *i*. In (19), we choose *z* to be equal to the number of silent sources, i.e., *z* = *k*, and *θ* is an exponential decay constant, which normalizes the distances of sources from each other in a discretized brain model, by their variance as follows:

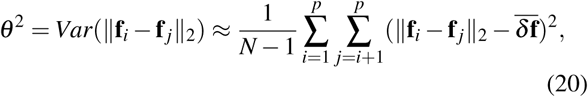

where 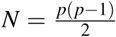 is the total number of inter-source distances, and 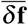 is an estimated average of these inter-source distances, given by:

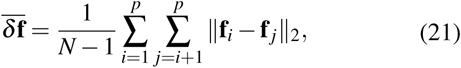

The degree matrix of the graph (**D**) is given by:

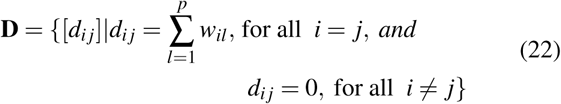

Using the degree and weight matrices defined in (19) and (22), the graph Laplacian matrix, **L** in (18), is defined as follows:

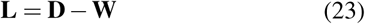

Based on one of the properties of the graph Laplacian matrix [41], we can write the quadratic term in the objective function of (18) as follows:

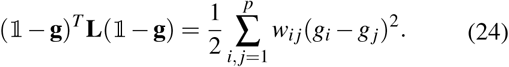

where **g** ∈ ℝ^*p*^. This quadratic term promotes the contiguity in the localized region of silence, e.g., an isolated node in the region of silence, which is surrounded by a number of active sources in the nearest neighbor graph, causes a large value in the quadratic term in (24), since *w*_*i j*_ has large value due to the contiguity, and the difference (*g*_*i*_ − *g* _*j*_) has large value, since it is evaluated between pairs of silent (small *g*_*i*_)-active (large *g* _*j*_) sources.

For a given *k*, the regularization parameter *λ* in (18), is found through a grid-search and the optimal value (*λ*^*^) is found as the one which minimizes the total normalized error of source contribution and the contiguity term as follows:

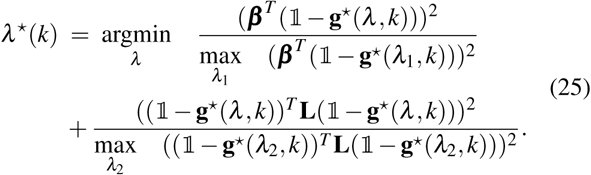

In addition, the size of region of silence, i.e., *k*, is estimated through a grid-search as follows:

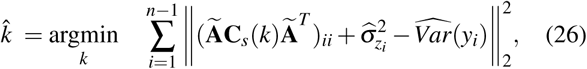

where (.)_*ii*_ indicates the element of a matrix at the intersection of the *i*^*th*^ row and the *i*^*th*^ column, 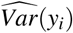 is the estimated variance of the *i*^*th*^ differential signal in **Y**, and 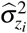 is the estimated noise variance at the *i*^*th*^ electrode location (see Appendix X B and X C to see how these might be estimated). In (26), *C*_*s*_(*k*) is the source covariance matrix, when there are *k* silent sources in the brain. The estimate 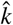 minimizes the cost function in (26), which is the squared error of difference between the powers of scalp differential signals, resulting from the region of silence with size *k*, and the estimated scalp powers based on the recorded data, with the measurement noise power removed. Under the assumption of identical distribution of sources, and lack of spatial correlation in the low-resolution source grid, and based on (10), we can rewrite (26) as follows:

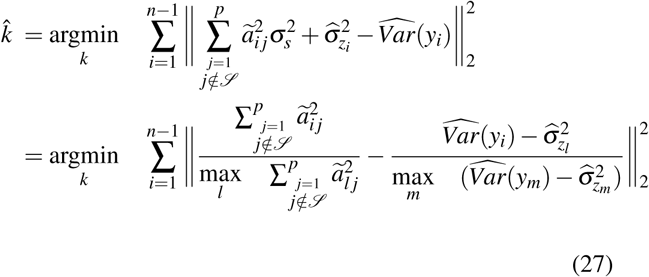

where *ã*_*i j*_ is the element of matrix **Ã** at the intersection of the *i*^*th*^ row and the *j*^*th*^ column, and 𝒮 is the set of indices of *k* silent sources, i.e., indices of sources corresponding to the *k* smallest values in **g**^*^(*λ*^*^, *k*), which is the solution of (18). The second equation in (27) normalizes the power of electrode *i* using the maximum power of scalp signals for each *i*. This step eliminates the need to estimate *σ*_*s*_ in the low-resolution.

Finally, the region of silence is estimated as the sources corresponding to the 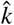 smallest values in 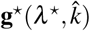. The 3D coordinates of the center-of-mass (COM) of the estimated contiguous region of silence in the low-resolution grid, i.e., 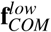, is used as an initial guess for the silence localization in the high-resolution grid, as explained in the next step.

#### Iterative algorithm based on a high-resolution grid and correlated sources

In this step, we use a high-resolution source grid, where the sources are not uncorrelated anymore. We try to estimate the source covariance matrix **C**_*s*_ based on the spatial exponential decay assumption in (9). In each iteration, based on the estimated source covariance matrix, the region of silence is localized using a CSpeC framework.

##### (i) Initialization

In this step, we initialize the source variance 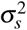, the exponential decay coefficient in the source co-variance matrix *γ*, and the set of indices of silent sources 𝒮 as follows:

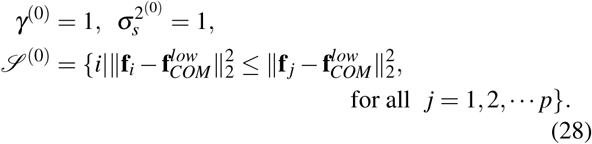

where 𝒮 ^(0)^ is simply the index of nearest source in the high-resolution grid to the COM of the localized region of silence in the low-resolution grid, i.e.,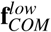.

For *r* = 1, 2, *…R*, we repeat the following steps until either the maximum number of iterations (*R*) is reached, or COM of the estimated region of silence 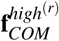 has converged, where 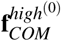 is the location of the source with index 𝒮 ^(0)^ in the high-resolution grid. The convergence criterion is defined as below:

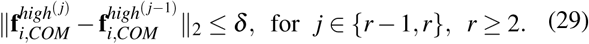

where *δ* is a convergence parameter for COM displacement through iterations, and 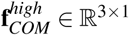.

##### (ii) Estimation of 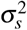 and γ

In this step, we estimate the source variance 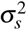, and the exponential decay coefficient of source covariance matrix *γ*, based on their values in the previous iteration and the indices of silent sources in 𝒮 ^(*r*−1)^. We define 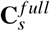 as the source co-variance matrix when there are no silent sources in the brain, and use it to measure the effect of region of silence on the power of each electrode. The source covariance matrix in the previous iteration (*r* − 1) is calculated as follows:

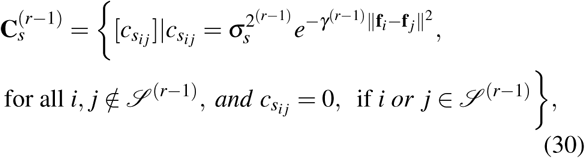

and 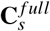 is given by:

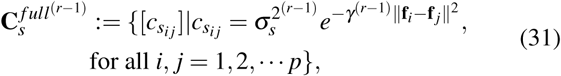

where there is no zero row and/or column, i.e., there is no silence. To be able to estimate 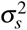 and *γ* based on the differentially recorded signals in **Y**, we need to find the electrodes which are the least affected by the region of silence. Based on the assumption (v) in Section IV A, the region of silence is much smaller than the non-silent brain region and some electrodes can be found on scalp which are not significantly affected by the region of silence. We find these electrodes by calculating a power-ratio for each electrode, i.e., the power of electrode when there is a silent region, divided by the power of electrode when there is not any region of silent in the brain, as follows:

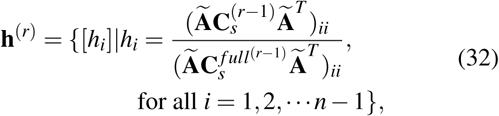

where **h** is a vector with values between 0 (all sources silent) and 1 (no silent source). Using this power ratio, we select the electrodes as follows:

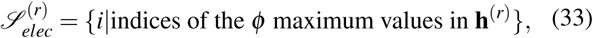

where 𝒮 _*elec*_ is the indices of the top *ϕ* electrodes which have the least power reduction due to the silent sources in 𝒮. Based on the differential signals of the selected *ϕ* electrodes in (33), *γ*^(*r*)^ and 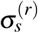 are estimated as the least-square solutions in the following equation:

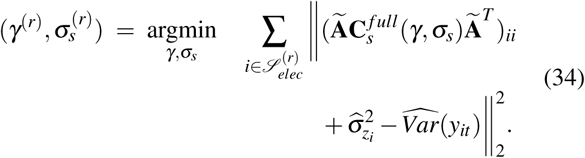

##### (iii) Localization of silent sources in the high-resolution grid

Based on the correlatedness assumption of sources in the high-resolution grid, we modify the source contribution measure definition (from equation (17)) as follows:

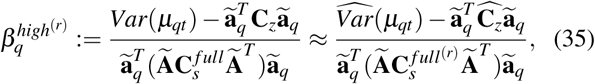

where 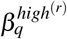 takes values between 0 (all sources silent), and 1 (no silent source in the brain). The only difference between 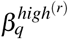 in the high-resolution grid and *β*_*q*_ in the low-resolution grid is in their denominators, which are essentially the variance terms in the absence of any silent source 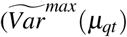 in (15)). In *β*_*q*_, the denominator is divided by the source variance 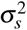, to be able to calculate *β*_*q*_ without estimation of 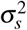. However, in the high-resolution grid, the denominator of 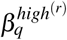 is simply 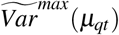, which is calculated under the source correlatedness assumption and using the estimated 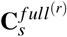. Using the definition of source contribution measure 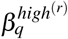 in the high-resolution grid, at iteration *r*, the contiguous region of silence is localized through a CSpeC framework, similar to the one defined in (18). However, we use the estimated source covariance matrix in each iteration to introduce a new set of constraints on the powers of the electrodes, which are less affected by the region of silence, i.e., the electrodes in 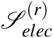, as defined in (33). Based on these power constraints, we obtain a convex optimization framework to localize the region of silence in the high-resolution brain model as follows:

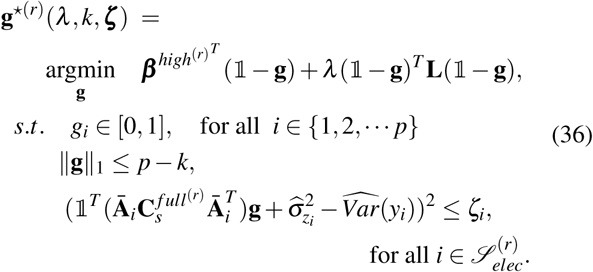

where 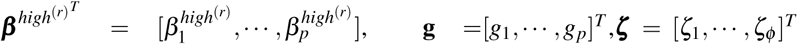, *λ* and *ζ*_*i*_ are regularization parameters, and **Ā**_*i*_ is a diagonal matrix, with the elements of *i*^*th*^ row of **Ã** on its main diagonal, defined as below:

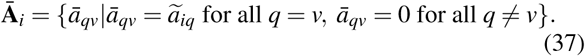

In (36), *ζ*_*i*_ is chosen to be equal to the square of the residual error in (34), for each *i* ∈ 𝒮_*elec*_, i.e.,

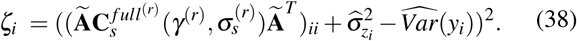

In each iteration *r*, values of *λ* and *k* are found in a similar way as they are found in the low-resolution grid (see equations (25) and (26)). However to estimate *k* based on (26), in the high-resolution grid we use 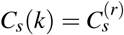, as is defined in (30). After each iteration, the set of silent indices in 𝒮 ^(*r*)^ is updated with the indices of the 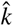 smallest values in the solution of (36), i.e., 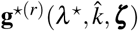.

After convergence, i.e., when the convergence criterion is met (see (29)), the final estimate of region of silence is the set of source indices in 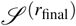.

#### Choosing the best reference electrode

the final solution 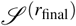 may change as we choose different EEG reference electrodes, which changes the matrix of differential signals of scalp **Y** and the forward matrix **Ã** in (7). The question is how to choose a reference electrode which gives us the best estimation of region of silence? To address this question, we use an approach similar to the estimation of 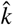, i.e., we choose the reference electrode which gives us the minimum scalp power mismatch. We define the power mismatch Δ*Pow* as follows:

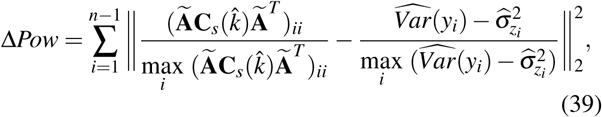

where both **Ã** and *y*_*i*_ are calculated based on a specific reference electrode. Δ*Pow* is the total squared error between the normalized powers of scalp differential signals, resulting from the region of silence with size 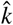, and the estimated scalp powers based on the recorded data with a specific reference.

### C. SilenceMap with baseline recordings

If we consider a hemispheric baseline or, more generally, have a baseline recording, the 2-step SilenceMap algorithm remains largely the same. In an ideal case where we have a baseline recording of scalp potentials, we simply compare the contribution of each source in the recorded scalp signals when there is a region of silence in the brain, with its contribution to the baseline recording. This results in a minor modification of the SilenceMap algorithm. The definitions of source contribution measures in (17) and (35), need to be changed as follows:

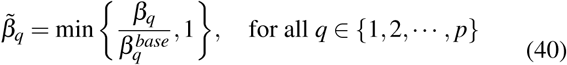

where *β*_*q*_ is defined in (17) for the low-resolution grid, and in (35) for the high-resolution grid, and 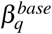 is the corresponding contribution measure of source *q* in the baseline recording. However, if the baseline recording is not available for the silence localization (as it was not available in our dataset used in the Results), based on the assumption of hemispheric symmetry in Section IV A (see assumption (viii)), one can use a hemispheric baseline. The source contribution measure is defined in a relative way, i.e., each source’s contribution measure is calculated in comparison with the corresponding source in the other hemisphere, as follows:

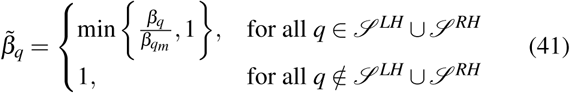

where 𝒮 ^*LH*^ is the set of indices of sources in the left hemisphere and 𝒮 ^*RH*^ is the set of indices of sources in the right hemisphere, and source indices which are not in 𝒮 ^*LH*^ ⋃ 𝒮 ^*RH*^, are located across the longitudinal fissure, which is defined as a strip of sources on the cortex, with a specific width *z*^*gap*^. The index *q*_*m*_ in (41) is the index of the “mirror” source for source *q*, i.e., source *q*’s corresponding source in the other hemisphere.

Equation (41) reveals the advantage of having a baseline for the silence localization task, i.e., we can relax the identical distribution assumption of sources in the source contribution measure, which makes 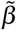 robust against the violation of the identical distribution assumption of sources in the real world. The rest of the algorithm remains the same, as is explained in Section IV B.

To find the solution of the CSpeC optimization in (18) and (36), *CVX*, a MATLAB package for specifying and solving convex programs [42, 43], is used. In addition, MATLAB nonlinear least-square solver is used to find the solution of (34).

### D. Modification of source localization algorithms to compare with the SilenceMap algorithm

To compare the performance of our SilenceMap algorithm with the state-of-the-art source localization algorithms, namely, MNE, MUSIC, and sLORETA, we modified them for the silence localization task. These modifications largely consist of adding additional steps to select the silent sources based on the estimated source localization in each algorithm. These modifications only make for a fairer analysis and answer the question of whether small modifications on existing source localization algorithms can localize silences. The details of these modifications are explained in this part.

#### Modified minimum norm estimation (MNE)

Minimum norm estimation (MNE) is one of the most commonly used source localization algorithms [14, 19]. In this algorithm, the brain source activities are estimated based on a minimal power assumption, and through the following regularization method:

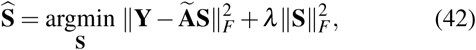

where **Ŝ** is the estimated matrix of source signals, **Ã** = **MA, Y** is the matrix of scalp differential signals defined in (6), *λ* is the regularization parameter, and ∥. ∥_*F*_ denotes the Frobenius norm of a matrix. Equation (42) has the following closed form solution:

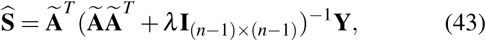

where **I** is the identity matrix, and *λ* is obtained using a grid-search and based on the *L* − *curve* method [44]. The MNE algorithm is kept unchanged until this point. **Ŝ**, the estimated localization across time, is used to localize silences. For a fair comparison, we do so by using the two step approach used in SilenceMap, i.e., we start from a low-resolution source grid and localize the region of silence, which is used as an initial guess for source localization in a high-resolution grid.

##### Low-resolution grid

In a low-resolution source grid, we localize the region of silence through the following steps: (i) We initialize the number of silent sources as 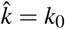; (ii) The squares of the elements in 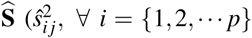, *∀ j* = *{*1, 2, *… t}*) are calculated for source power comparison; (iii) For each time point *j*, sort the estimated source powers 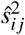 in the ascending order and choose the first 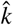 corresponding sources, which are the sources with the minimum power at time *j*. We name the set of indices of these sources as 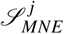; (iv) Based on the repetition of sources in 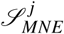, we calculate a histogram (*hist* _*MNE*_). Then this histogram is normalized and sorted in the descending order (the source with the largest population of 1 has the first index). The normalized population of source *q* is shown as 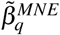 ; (v) In this step, we find an estimate of the size of region of silence 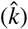. This is done by finding the knee point in the curve of 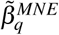 vs. *q* (see Fig. 6a). In a curve, the “knee” point is defined as the point where the curve has maximum curvature, i.e., the point where the curve is significantly different from a straight line [45–47]. To find the knee point in the curve of 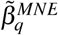 vs. *q*, we define a measure of distance to the origin (*q* = 0, 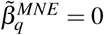) as follows:

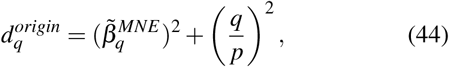

where 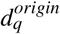 is the defined distance of point (*q* = 0, 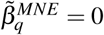) on the curve to the origin, and *p* is the total number of sources in the descritized brain model. Fig. 6b shows the calculated 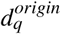 for the curve in Fig. 6a. We choose the closest point to the origin as the knee point 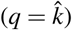, where the index of this knee point 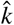 is an estimation for *k* (see Fig. 6b).

**FIG. 6.**
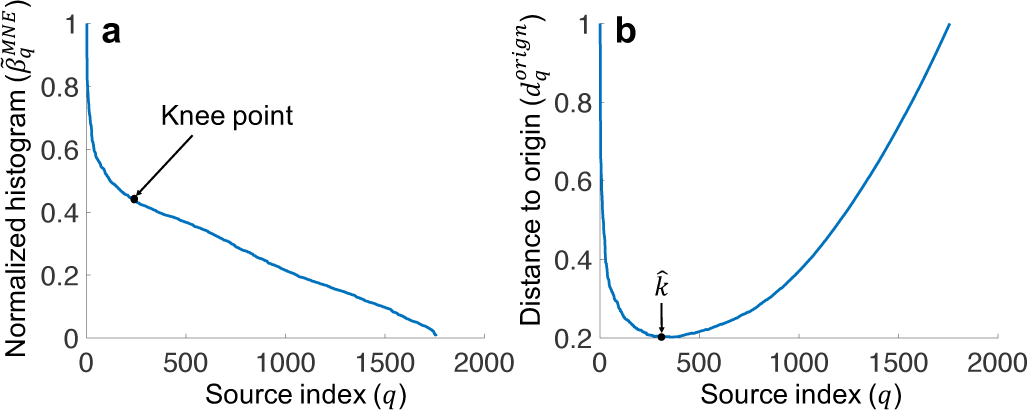
Estimation of the size of the region of silence (*k*) in the modified MNE algorithm based on the knee point detection: a) 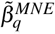 is the normalized and sorted histogram of sources in a descending order, which captures the frequency of a source to being among the 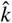 sources with the minimum power over time. In this curve, a “Knee” point is defined as the point with maximum curvature, i.e., the point where the curve is significantly different from a straight line; b) Distances of points on the curve in (a) from the origin (*q* = 0, 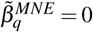). The index of the point with the minimum distance from the origin (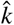) is chosen as an estimation of *k*.

(vi) In this step, we exploit the knowledge of contiguity of the region of silence and estimate the region based on the estimated number of silent sources 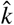. First we choose the 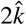 sources with the minimum power over time, i.e., the 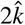 sources which have maximum 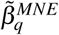. Then we calculate the COM of the 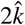 selected sources in the low-resolution grid 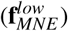, and choose the 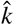-nearest neighbors of 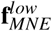 as the estimated region of silence in the low-resolution grid.

##### High-resolution grid

We use the COM of the estimated region of silence in the low-resolution grid 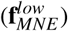, as an initial guess and try to improve the localization performance in a high-resolution source grid. The steps are mainly the same as the steps used in the low-resolution grid, except in the last step (vi), where we use the COM of the estimated region of silence in the low-resolution grid, and choose the 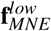-nearest neighbors of 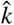 as the estimated region of silence in the high-resolution grid, where 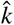 is the estimated size of region of silence in the high-resolution grid based on the knee point detection method in step (v).

#### Modified multiple signal classification (MUSIC)

Multiple signal classification (MUSIC) is a source localization algorithm, which is based on a sequential search of sources, rather than finding all sources at the same time [17, 18]. In MUSIC, the singular value decomposition (SVD) of the matrix of scalp recording signals 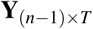 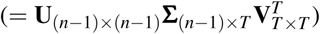 is used to reconstruct an orthogonal projection to the noise space of **Y** to quantify the contribution of each source in the recorded signal **Y** [14]. The MUSIC algorithm follows these steps for source localization: (i) We select the left singular vectors (columns of **U**) which correspond to the large singular values up to *ρ*% of the total energy of the matrix 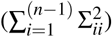, where *ρ* is a constant. These selected singular vectors (**U**_*s*_) form a basis for the observation data; (ii) We construct an orthogonal projection matrix to the noise space of **Y** as 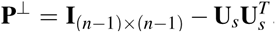. Using this matrix the MUSIC cost function is written as [14]:

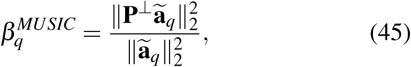

where **ã**_*q*_ is the *q*^*th*^ columns in **Ã**, and 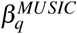 is a measure of contribution of source *q* in the noise space of the recorded scalp potentials in **Y**. The MUSIC algorithm is kept unchanged until this point. The next steps of the modified MUSIC algorithm, in both low-resolution and high-resolution grids closely follow the last two steps in the Modified MNE algorithm, and we use the measure of source contribution in MUSIC 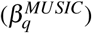 instead of 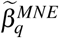, where the source with no contribution in the differential measured signal **Y** has 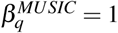. Therefore, the main difference between the MUSIC algorithm and the modified MUSIC, is that in the MUSIC, the measure of contribution of source 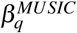 is used to fined the active sources, i.e., the sources with small 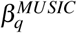 values, while for the silence localization the sources with large 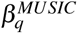 values are selected based on the contiguity assumption of the region of silence and using the knee point thresholding mechanism (step (v) in the modified MNE).

#### Modified standardized low resolution brain electromagnetic tomography (sLORETA)

We modify the source localization sLORETA algorithm, introduced in [21], in the same way that we modified the MNE algorithms for the silence localization. However, the minimum-norm solution requires an additional step of normalization by the estimated source variances. Since we assume that the orientations of dipoles in the brain are known, i.e., they are normal to the surface of the brain, following equation (22) in [21], the estimated power of source activities in the brain based on the sLORETA algorithm is given by the following equation:

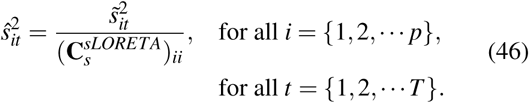

where 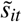 is the *i*^*th*^ element of the *t*^*th*^ column in the minimum-norm solution 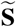, which is given by equation (43), and 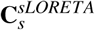 is defined as [21]:

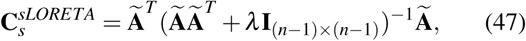

**Ŝ** in (46) is used for the silence localization task, following the steps mentioned for the modified MNE algorithm. The minimum-norm solution in the sLORETA algorithm, as is defined in [21], is based on an average reference for the scalp potentials. However, we rewrite the minimum-norm solution as 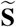 in (43) based on a specific reference electrode, rather than the average reference electrode. The parameters used in the implementation of these modified source localization algorithms are available in the Appendix X D.

### E. Data analysis

#### Prepossessing steps

We preprocess the recorded EEG signals using *EEGLAB* [48] toolbox in MATLAB. First, we bandpass filter the EEG data in the frequency range of [1, 100] *Hz* using a Hamming windowed sinc finite impulse response (FIR) filter. Then, we visually inspect the noisy channels, remove and spatially interpolate them. In the next step, we calculate two differential channels based on the pairs of eye electrodes, one for vertical and one for the horizontal eye movements, and along with the heart channel and all scalp electrodes, an independent component analysis (ICA) is applied to remove the eye artifacts and heart beats from the EEG signals. After removing the artifact components from the EEG signals, we examine the channels one more time using the channel statistics, where a normal distribution is fitted to the data of each channel and based on the standard deviation, skewness, and kurtosis, channels with significantly different statistics are removed and interpolated. Finally, the signals are epoched into 2-second intervals and epochs with abnormal trends, values, and/or abnormal power spectral densities are removed, using the *EEGLAB* toolbox.

#### Ground truth regions and MRI scans

The ground truth regions of silence are extracted based on the MRI scans of patients (see Fig. 2) following these steps: (i) 3D models of descritized cortex are extracted by processing the MRI scans using the *FreeSurfer* software [49–55], and removing the layers of the head, namely, CSF, skull, and scalp using the *MNE* open-source software [56], (ii) the sources/nodes of the intact hemisphere, i.e., the hemisphere without any missing part, are mirrored along the longitudinal fissure, (iii) the smallest distance of the mirrored sources are calculated from the sources in the hemisphere with the resected part, (iv) *N* sources with the largest distance are selected as the region of silence, where *N* is determined by visual comparison of the extracted ground truth in the 3D model, and its corresponding MRI scan.

Displayed figures in this paper are generated using MATLAB, Microsoft PowerPoint, and *FreeSurfer* software.

### F. Simulated Dataset

We simulate EEG signals at 128 electrodes, located at the 10-5 standard system of scalp locations [30], as follows: (i) First, we use a high-density source grid, extracted by discretizing a real brain model, and randomly choose a node along with its *k* nearest neighbors, as the region of silence with size *k*, (ii) then we simulate the source signals using a multi-variate gaussian random process with a covariance matrix **C**_*s*_ defined in (9), where 𝒮 is the set of indices of silent sources specified in the first step, the source variance is *σ*_*s*_ = 1 *mV*, and the exponential decay coefficient is *γ* = 0.12 (*mm*)^−2^. (iii) the measurement noise in (7), i.e., 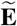, is simulated using a multivariate gaussian random process with a covariance matrix **C**_*z*_ defined in (8), where 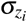 is chosen randomly from a uniform distribution in the range of 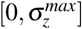, and 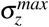 is chosen so that the baseline EEG signals, i.e., without any region of silence, have a specific average SNR, defined as below [57–59]:

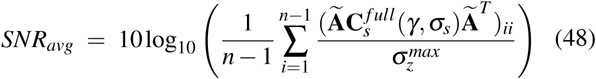

(iv) In the next step, the forward matrix **A** in (5) is calculated based on a real head model, which is obtained from MRI scan of patient OT in the real database (Fig. 2). In this paper, we use the *FreeSurfer* software [49–55] to process the MRI images, and use *MNE* open-source software [56] to extract different layers of the head, i.e., brain, cerebrospinal fluid (CSF), skull, and scalp. Then having these layers of head, and using the boundary element method (BEM), the forward matrix **A** is computed, where a free MATLAB toolbox, *Field-Trip* [60], is used. In the BEM model, volume conductivity ratios of [1, 0.067, 5, 1] are used for scalp, skull [61], CSF, and brain respectively. We take the intact hemisphere (the one without any missing parts) and mirror it across the longitudinal fissure to form a symmetric brain model, which is used as the source grid in our algorithm. In Section III we discuss and quantify the localization error we introduce by using this symmetric brain model.

## V. DATA AVAILABILITY

The anonymized raw EEG dataset and MRI scans (shown in Fig.2) of the participants in this research are made available online on KiltHub, Carnegie Mellon University’s online data repository (DOI: 10.1184/R1/12402416).

## VI. CODE AVAILABILITY

The SilenceMap algorithm was developed in MATLAB, using standard toolboxes, and the *CVX* MATLAB package [42, 43]. All MATLAB code is made available online on GitHub (DOI: 10.5281/zenodo.3892185).

## VIII. ACKNOWLEDGMENTS

This work was supported in part by the Chuck Noll Foundation for Brain Injury Research, and a CMU BrainHub award. The authors of this manuscript thank Chaitanya Goswami, Praveen Venkatesh, Ashwati Krishnan, Anne Margarette S. Maallo, Sarah M. Haigh, Animesh Kumar, and Maysamreza Chamanzar for helpful discussions, and Ashwati Krishnan and Patricia Brosseau for help in data collection.

## IX. AUTHOR CONTRIBUTIONS

A.C., M.B., and P.G. initiated, designed, and executed the research. A.C. and M.B. acquired and interpreted the data. M.B. and P.G. supervised the data collection. A.C. and P.G. developed the algorithms. A.C. developed the software tools necessary for conducting the experiments and analyzing the data. A.C., M.B., and P.G. wrote the manuscript.

## X. SUPPLEMENTARY MATERIALS

### A. Estimation of sample variance of *µ*_*qt*_

To estimate *Var*(*µ*_*qt*_) in (13), average of the sample variances of *µ*_*qt*_ cannot be used since based on the *WSS* assumptions in Section IV A, the elements of **S** and 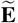, and consequently *µ*_*qt*_ are correlated over time. However, since *µ*_*qt*_ in (12) is a linear combination of *s*_*it*_ and 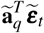, which are both *WSS* and are not correlated with each other, *µ*_*qt*_ is also *WSS* and its variance can be estimated numerically using the time samples, as follows [62]:

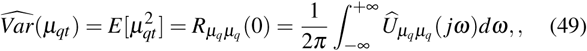

where 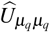 is an estimation of power spectral density (PSD) of correlation coefficient *µ*_*qt*_. We have used Welch’s method to obtain this estimate [63–65], which is implemented in Matlab. A window size of 500ms, with 250ms overlap used in estimation of the PSD.

### B. Estimation of noise covariance matrix C_*z*_

As mentioned in Section I, the main difference between source localization and silence localization is in the noise definition. Most of the source localization algorithms group together the measurement noise with the background brain activity and provide methods for estimation of noise [12, 13]. However, the background brain activity is crucial for the silence localization to distinguish between normal brain activity and abnormal silence. Therefore we need to revise the noise definition accordingly. As mentioned in Section IV A, noise is white and bandpass filtered in a specific frequency interval of [*f*_*L*_ = 1, *f*_*H*_ = 100]*Hz* (during the preprocessing step, see Section IV E). *f*_*L*_ and *f*_*H*_ are the lower and the upper cutoff frequencies of the filter we have used in the preprocessing step to bandpass filter the scalp EEG signals. In addition, we assume that the noise components are spatially uncorrelated, and the noise covariance matrix is a diagonal matrix, as is defined in (8). Therefore, under stationary assumption for 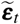, we can estimate 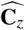 as follows:

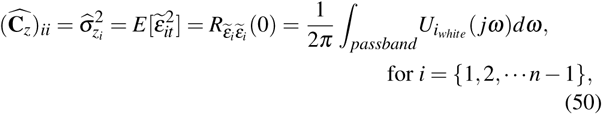

where 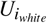 (*jω*) is the constant power spectral density of the white noise at the scalp electrode *i* in the frequency bands of [*f*_*L*_, *f*_*H*_] and [− *f*_*H*_, − *f*_*L*_], and zero outside these frequency intervals, and 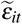 is the element of 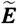 at *t*^*th*^ time point and *i* ^*th*^ electrode (see equation (7)). This constant power spectral density can be estimated from the power spectral density of the recorded signal *y*_*i,t*_ in the high fre quencies (≥ *f*_*H*_ − Δ *f*), where we assume that EEG does not have any frequency component and the noise power dominates:

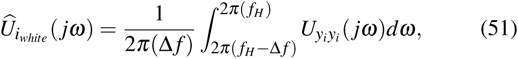

where Δ *f* is the frequency bandwidth in which the white noise is assumed to have the dominant energy (see Fig. 7).

**FIG. 7.**
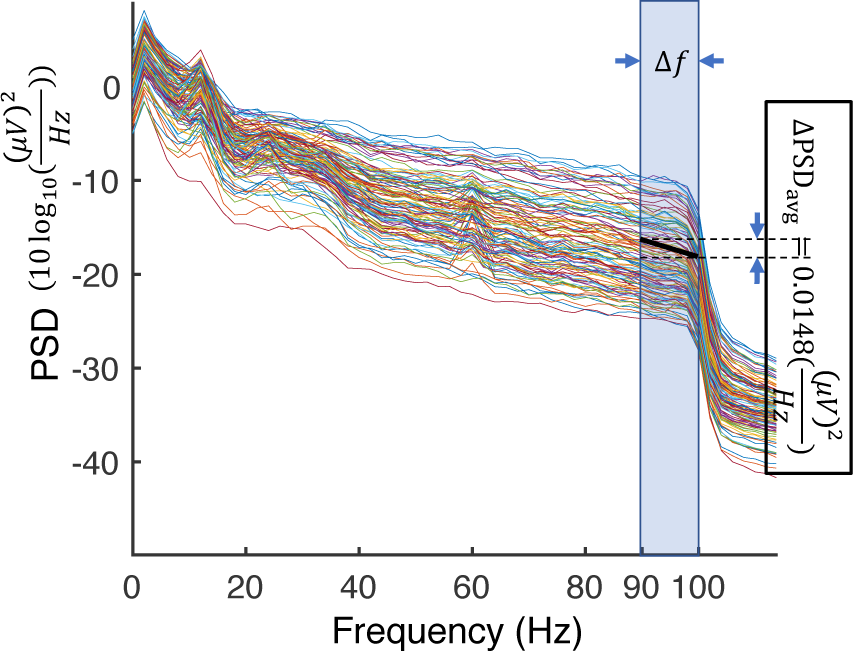
PSD of differential signals in a healthy subject (DH). The signals are bandpassed in the frequency interval of [*f*_*L*_ = 1, *f*_*H*_ = 100]*Hz*. The highlighted region is the frequency interval, during which the approximately white noise is assumed to have the dominant energy. On average, the PSD of EEG signals drops only by 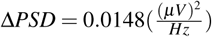 from 90*Hz* to 100*Hz*.

where 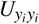 (*jω*) can be estimated from the differential signal in the *i*^*th*^ row of **Y** (see Section X C). Modeling and estimation of EEG noise based on the PSD of the recorded signals is commonly used in EEG source localization studies [66–68], which we have modified for the silence localization task. Fig. 7 shows the PSD of the differential signals in a healthy subject (DH), which are bandpassed in the frequency interval of [*f*_*L*_ = 1, *f*_*H*_ = 100]*Hz* during the preprocessing step. The frequency interval, during which white noise is assumed to have the dominant energy is highlighted in this figure (Δ *f* = 10*Hz*). This assumption is approximately true, i.e., averaged over *i* = 1, 2,*… n* − 1, the PSD of *y*_*i*_ drops only by 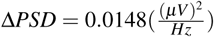 from *f*_*H*_ − Δ *f* = 90*Hz* to *f*_*H*_ = 100*Hz* (see Fig. 7).

### C. Estimation of the variance of differential signals in Y

Similar to the estimation of the noise variance in Section X B, we estimate 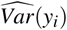 based on its PSD. An average of the sample variances of *y*_*i*_ cannot be used since based on the *WSS* assumptions in Section IV A, the elements of **S** and 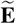, and consequently *y*_*i*_ are correlated over time. However, since *y*_*i*_ in (11) is a linear combination of *s*_*it*_ and 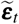, which are both *WSS* and are not correlated with each other, *y*_*i*_ is also *WSS* and its variance can be estimated numerically using the time samples, as follows [62]:

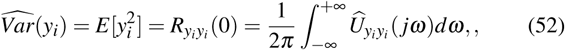

where 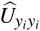 is an estimation of PSD of the differential signal *y*_*i*_. We have used Welch’s method to obtain this estimation [63–65].

### D. The list of all parameters and their values used in the SilenceMap algorithm and modified source localization algorithms

The values of all parameters we have used to implement and test the SilenceMap algorithm, with and without baseline, as well as the modified source localization algorithms are summarized in Table III. All code and datasets are freely available online in [69, 70].

**TABLE III.**
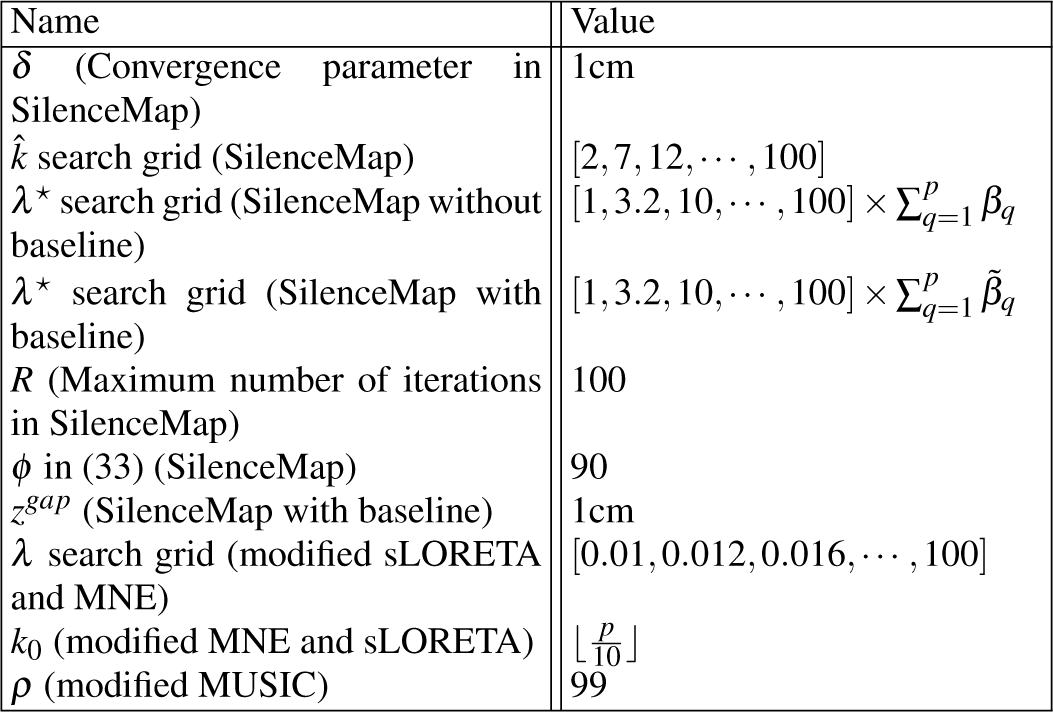
Parameters used for implementation of the SilenceMap, modified MNE, MUSIC, and sLORETA algorithm.

| Name | Value |
| --- | --- |
| $\delta$ (Convergence parameter in SilenceMap) | 1cm |
| $\hat{k}$ search grid (SilenceMap) | $[2, 7, 12, \dots, 100]$ |
| $\lambda^*$ search grid (SilenceMap without baseline) | $[1, 3.2, 10, \dots, 100] \times \sum_{q=1}^p \beta_q$ |
| $\lambda^*$ search grid (SilenceMap with baseline) | $[1, 3.2, 10, \dots, 100] \times \sum_{q=1}^p \tilde{\beta}_q$ |
| $R$ (Maximum number of iterations in SilenceMap) | 100 |
| $\phi$ in (33) (SilenceMap) | 90 |
| $z^{gap}$ (SilenceMap with baseline) | 1cm |
| $\lambda$ search grid (modified sLORETA and MNE) | $[0.01, 0.012, 0.016, \dots, 100]$ |
| $k_0$ (modified MNE and sLORETA) | $\lfloor \frac{p}{10} \rfloor$ |
| $\rho$ (modified MUSIC) | 99 |

## Footnotes

1 E.g. authors in [12] state: “EEG data are always contaminated by noise e.g., exogenous noise and background brain activity”.

